# Cardiac-locked auditory stimulation modulates pupil and neural dynamics

**DOI:** 10.1101/2025.06.26.661871

**Authors:** Giovanni Chiarion, Jacinthe Cataldi, Sophie Schwartz, Andria Pelentritou, Marzia De Lucia

**Affiliations:** Brain-Body and Consciousness Laboratory, Department of Clinical Neurosciences, Lausanne University Hospital, University of Lausanne, 1011 Lausanne, Switzerland; Centre for Biomedical Imaging, 1011 Lausanne, Switzerland; Department of Neuroscience, Faculty of Medicine, University of Geneva, 1211 Geneva, Switzerland; Swiss Center for Affective Sciences, University of Geneva, 1202 Geneva, Switzerland

**Keywords:** auditory regularity, pupil diameter, global field power, heartbeat, brain-body interactions, auditory evoked potential, cardio-audio, EEG

## Abstract

The human brain is sensitive to temporal regularities across bodily and environmental signals. Here, we investigated the pupil and neural correlates of regularity encoding established across cardiac and auditory stimuli. Auditory sequences were presented in synchrony with the heartbeat (synchronous), at a fixed pace, or without temporal regularity while recording pupillometry, electroencephalography, and electrocardiography in healthy participants. Sounds evoked typical pupil dilation in all conditions. However, only in the synchronous condition, pupil dilation progressively decreased over the course of the sequence, possibly reflecting adaptation to the repeated cardio-auditory alignment. A concurrent increase in global EEG activity suggested enhanced cortical processing in response to the synchronous sequence. Pupil constriction was driven by participants with higher heart rate, indicating that pupil adaptation mostly occurs in response to fast auditory sequences. Cardio-audio regularity encoding manifests as a pupil adaptation and an amplification of global EEG activity, likely reflecting improved temporal prediction precision.

## Introduction

Identifying patterns in the sensory environment is a fundamental function of the human brain, supporting prediction^1^, shaping perception^2^ and guiding decision-making^3^. Research shows that humans can detect regular patterns in sound sequences, often implicitly and effortlessly^4^. The neural and peripheral mechanisms underlying auditory regularity encoding have been commonly investigated using paradigms that introduce deviant sounds or sound omission within the ongoing regular auditory sequence^5^. Such violations of regularity can trigger neural and physiological responses which provide markers of successful regularity encoding, and are associated with surprise^6^, attention reorientation^7^, and internal model updating^8^. One key physiological marker is pupil dilation, mostly under the control of the neuromodulator norepinephrine (NE) generated in the brainstem’s locus coeruleus (LC), with extensive projections to the brain and spinal cord^9^. Pupil dilation increases following the unexpected violation of regular patterns in auditory and visual sequences^10–13^, covaries with LC activation^14,15^ and is typically interpreted as a marker of arousal and cognitive load^16^. At the neural level, sensitivity to sensory regularities is commonly assessed using mismatch negativity, a well-established electroencephalographic (EEG) marker of auditory deviance detection^17^. In addition, encoding of auditory regularities is reflected in the increase of sustained global evoked activity such as the global field power (GFP), in response to regular patterns compared to random auditory sequences^18–21^.

Beyond external sensory input, recent studies highlight the key role of bodily signals in modulating auditory regularity processing^22^. In particular, integration of cardiac and auditory stimuli was shown by studies investigating the encoding of cardiac-based auditory regularity during wakefulness, sleep and coma^23–26^ and conscious processing of narrative stimuli in relation to cardiac modulation^27^.

An open question arising from previous research is whether temporal regularities driven by cardiac signals, engage both neural and autonomic systems - as reflected in EEG and pupil responses -, and whether these responses are coordinated and differ from those elicited by purely external temporal regularities. We therefore investigated the pupil and neural correlates of regularity encoding while administering auditory sequences under three distinct conditions in healthy volunteers: synchronous (synch), in which each sound onset was temporally locked to the individual’s electrocardiography (ECG) R-peak; isochronous (isoch), where sounds were presented at fixed, predictable intervals (between successive sounds); and asynchronous (asynch), featuring irregular intervals with no relationship to the cardiac rhythm. We hypothesized that both the synch and isoch conditions - each enabling the formation of auditory expectations, either through heartbeat coupling (synch) or fixed sound-to-sound (SS) intervals (isoch) - would elicit distinct autonomic and neural responses compared to the asynch condition, which lacks both forms of temporal predictability. Specifically, we predicted a gradual reduction in pupil diameter throughout the synch and isoch sequences relative to the sequence onset, reflecting LC adaptation and tonic arousal following successful regularity encoding^11,12^. At the neural activity level, in the synch condition, we expected a progressive increase in evoked activity from the onset of the auditory sequence, mirroring the formation of a cardio-auditory associative trace as previously described for other types of regularity encoding^18,19^. We anticipated this increase to be slower in the synch condition compared to the isoch, due to the additional complexity of integrating cardiac and auditory stimuli. Finally, we investigated whether the known relationship between pupil size and physiological signals such as the heart rate^27,28^ would hold, especially when the cardiac signals predict sound occurrence (synch condition). We thus hypothesized that pupil dynamics would interact with the ongoing cardiac rhythm, specifically in the synch condition, due to the bidirectional influence of sound predictability and heart rate in this condition^24,26^.

## Results

We analyzed the normalized pupil diameter and concurrent EEG activity in 31 healthy participants. The experiment consisted of six blocks, each containing the four conditions (no auditory stimulation, synch, asynch, isoch). Here, we report the results based on the data collected during the first block of the experiment. This block provided sufficient data to assess effects over the course of each auditory sequence of 300 stimuli, while minimizing the impact of fatigue in this passive listening experiment. The duration of this first block was comparable to previous experiments examining pupil responses to auditory regularities^29,7,11,12^. For completeness, results obtained from all six experimental blocks are provided in the Supplemental Information (Figures S1-S5). While similar trends were observed across blocks, many effects did not reach significance, possibly due to increased fatigue and adaptation during the two-to three-hour experimental session.

Participants passively listened to auditory sequences designed to probe responses to different temporal regularities (Figure 1): a cardio-audio regular sequence (synch), an isochronous sequence (isoch), and a sequence without regularity (asynch). To control for potential temporal expectation effects inherent in the structured timing of these auditory conditions, we generated three corresponding baselines from a fourth condition without auditory stimulation. Specifically, we retrospectively defined ‘artificial’ sound onsets in the data acquired without auditory stimulation by utilizing the sound onsets extracted from each auditory condition. For the synch baseline, we extracted pupil traces at 52 ms after each detected R-peak, matching the actual synch condition timing. For the isoch and asynch baseline, we segmented the pupil data based on the SS intervals from the corresponding auditory conditions. This approach enabled the direct comparison of evoked pupil responses while controlling for stimulus timing effects, since any observed differences could be attributed to the actual presence of auditory stimuli rather than the temporal structure of stimulus presentation. We examined pupil responses time-locked to both actual (synch, asynch, isoch) and ‘artificial’ (baselines) sound onsets. Our investigation focused on two distinct temporal scales: (i) local effects, assessing pupil and neural responses to individual sounds and, (ii) global effects, capturing cumulative changes in pupil diameter and neural activity over the course of each auditory sequence. Finally, we explored whether global pupil responses were related to the ongoing cardiac rhythm across the different experimental conditions.

**Figure 1.**
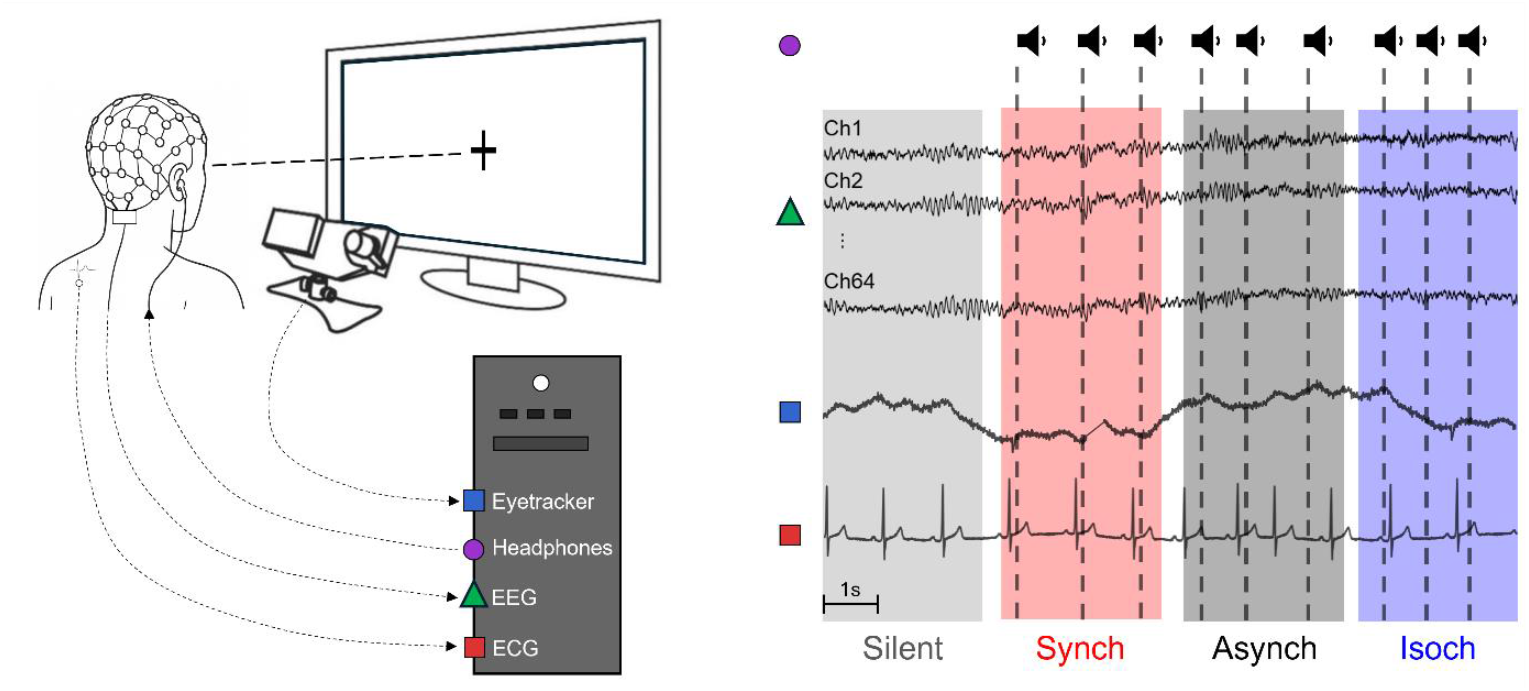
Data acquisition and auditory stimulation paradigm. Continuous EEG, ECG and pupil diameter recordings were acquired while participants fixated on a central cross and passively listened to sequences of pure tones containing different temporal regularities. The left panel illustrates the experimental setup, including the simultaneous recording of neural activity (64-channel EEG), cardiac activity (ECG), and pupil responses (eye-tracking). The right panel displays the four experimental conditions. In the Silent condition (gray), no auditory stimuli were presented; in the Synch condition (red), sounds were administered at a fixed delay following the real-time detection of R-peaks in the ECG; in the Asynch condition (black), sounds occurred with variable R peak-to-sound and sound-to-sound intervals, with no systematic relationship to the heartbeat; in the Isoch condition (blue), sounds were presented at a fixed sound-to-sound interval. Dashed vertical lines mark sound onsets across conditions.

### Pupil response to sounds

To assess local, sound-evoked pupil responses, for each participant (N = 26), we analyzed the normalized pupil diameter time-locked to sound onsets for the three auditory conditions, and to the ‘artificial’ sound onsets for the corresponding baselines. Here, each pupil response was referenced to the pre-stimulus period of 100 ms. Qualitatively, we observed that across all three auditory conditions (synch, asynch, isoch), sounds elicited a characteristic biphasic pupil response: an initial constriction followed by a dilation peaking around 500 ms after stimulus onset Figure 2a). By contrast, when no auditory stimuli were presented, the pupil response did not exhibit such a biphasic pattern (Figure 2a). To quantify the pupil diameter in response to sound vs baseline, pupil diameter was averaged over the entire trial window, yielding a single pupil response value per condition and participant. Comparisons between each auditory condition and its corresponding baseline using one-tailed Wilcoxon signed-rank tests (p < 0.05) revealed significantly higher pupil dilation in the synch (W = 88, p = 0.01), asynch (W = 63, p < 0.01), and isoch (W = 99, p = 0.02) conditions (Figure 2a, inset). A Friedman test contrasting the pupil response between the three auditory conditions (synch, asynch, isoch) revealed no significant differences (W < 0.01, p = 0.89), suggesting that while auditory stimuli reliably elicited a pupil response relative to baseline, the specific temporal structure of the sound sequences did not modulate the magnitude of the local pupil dilation.

**Figure 2.**
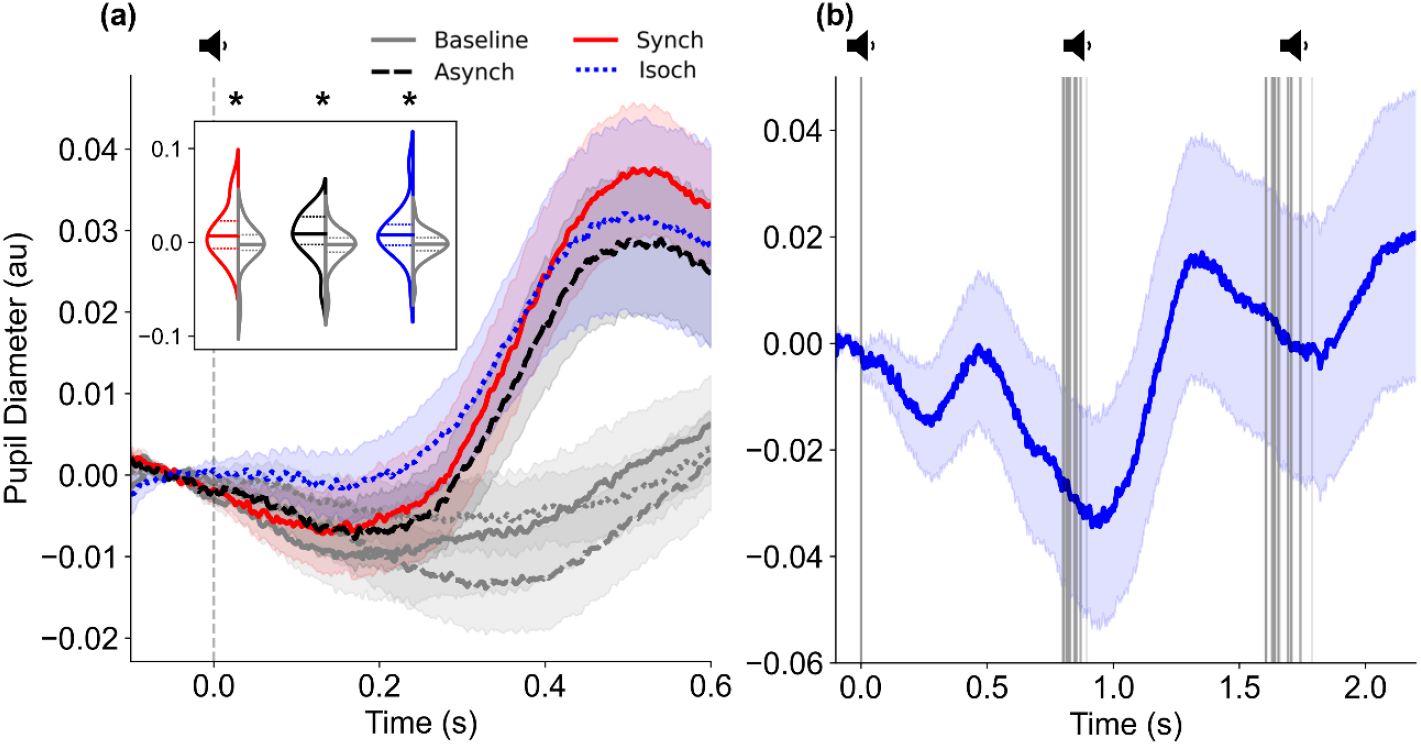
Pupil responses to sounds. (a) Grand-averaged pupil diameter (N = 26) time-locked to sound onset (vertical dashed gray line) across all conditions: Synch (red), Asynch (black), Isoch (blue), and their corresponding Baselines (gray). The Baseline responses represent pupil activity without sound presentation, epoched relative to ‘artificial’ sound onsets that matched the temporal structure of the auditory conditions. Pupil diameter data were z-score normalized and referenced to the median pupil diameter during the 100 ms pre-stimulus period. The inset shows violin plots of the averaged pupil diameter over the entire trial period (between -100 ms and 600 ms) for each auditory condition and its corresponding baseline. Horizontal lines within each violin plot indicate the first quartile, median, and third quartile. (b) Grand-averaged pupil diameter (N = 12) in response to sequences of three consecutive sounds (vertical gray lines) in the Isoch condition, time-locked to the first sound. Data were z-score normalized and referenced to the median pupil diameter during the 100 ms pre-stimulus period of the first sound. Shaded areas in both panels represent ± standard error of the mean. au = arbitrary units; *p < 0.05 by one-tailed Wilcoxon signed-rank tests.

Next, we investigated whether rapid sound presentation within the sequences could induce distinct pupil responses at each sound onset, despite their close temporal spacing. This analysis was performed only for the isoch condition, where fixed SS intervals allowed for the extraction of trials containing three consecutive sounds in a subset of participants with similar SS intervals (N = 12). Grand-averaged pupil responses across these triplets of pupil diameter changes were sufficiently rapid to be modulated by single sounds, with a local dilation after each auditory stimulus (Figure 2b).

### Neural response to sounds

To characterize neural responses to auditory stimuli with different temporal regularities, we computed the EEG auditory evoked potentials (AEPs), time-locked to sound onset (Figure 3a). While the asynch and isoch conditions yielded very similar AEP waveform morphologies and amplitudes, the synch condition revealed a distinct neural response (Figure 3b). Specifically, the synch condition showed a unique peak around 50 ms before sound onset, corresponding to the neural response to the heartbeat, since auditory stimuli in this condition were time-locked to heartbeats. Despite this difference, all three auditory conditions elicited a clear N100 component at approximately 100 ms post-stimulus onset with the expected spatial distribution characterized by maximal negative voltage amplitudes over central electrodes. GFP analysis confirmed the AEP findings, showing peak activity at approximately 100 ms post-stimulus onset across all conditions, consistent with the N100 component timing observed in the AEPs. The asynch and isoch conditions again displayed comparable GFP amplitudes and temporal dynamics. In the synch condition however, the GFP signal reflected a more complex pattern, likely resulting from the combined influence of cardiac-related activity preceding sound onset and the subsequent auditory response.

**Figure 3.**
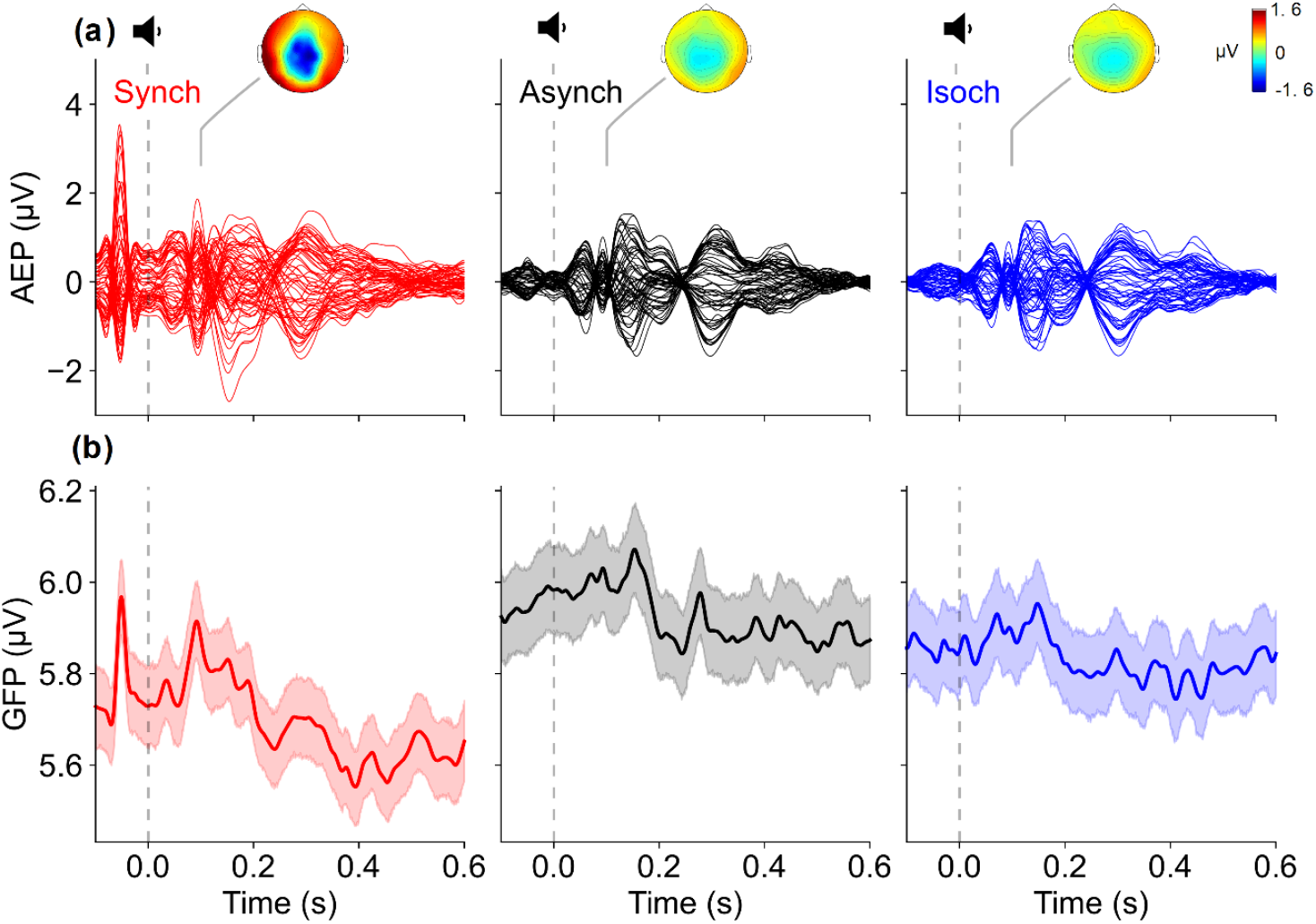
Neural auditory evoked responses. (a) Grand-averaged auditory evoked potentials (AEPs; N = 31), time-locked to sound onset (vertical dashed gray line) across the three auditory conditions: Synch (red), Asynch (black), and Isoch (blue). The topographic maps show the scalp distribution of AEP amplitudes at 100 ms post-stimulus onset. (b) Grand-averaged global field power (GFP) for each auditory condition, representing the overall strength of neural activity across all electrodes over time. GFP was computed as the standard deviation of EEG potentials across all electrodes at each time point, providing a reference-independent measure of global brain activity. Shaded regions represent ± standard error of the mean.

### Pupil and neural responses to sound sequences

To assess cumulative pupil changes during repeated auditory stimulation, we analyzed pupil responses across the full sequence of 300 sounds in each condition (N = 25). For each sound, the pupil diameter was averaged over the entire trial length (between -100 ms and 600 ms relative to each sound onset) and referenced to the mean pupil diameter from the first 10 trials of the same condition. This approach yielded a global pupil response trend across the entire sequence under different temporal regularities, capturing a systematic change in pupil size - either constriction (negative pupil diameter) or dilation (positive pupil diameter) - relative to the beginning of the auditory stimulation (Figure 4a).

**Figure 4.**
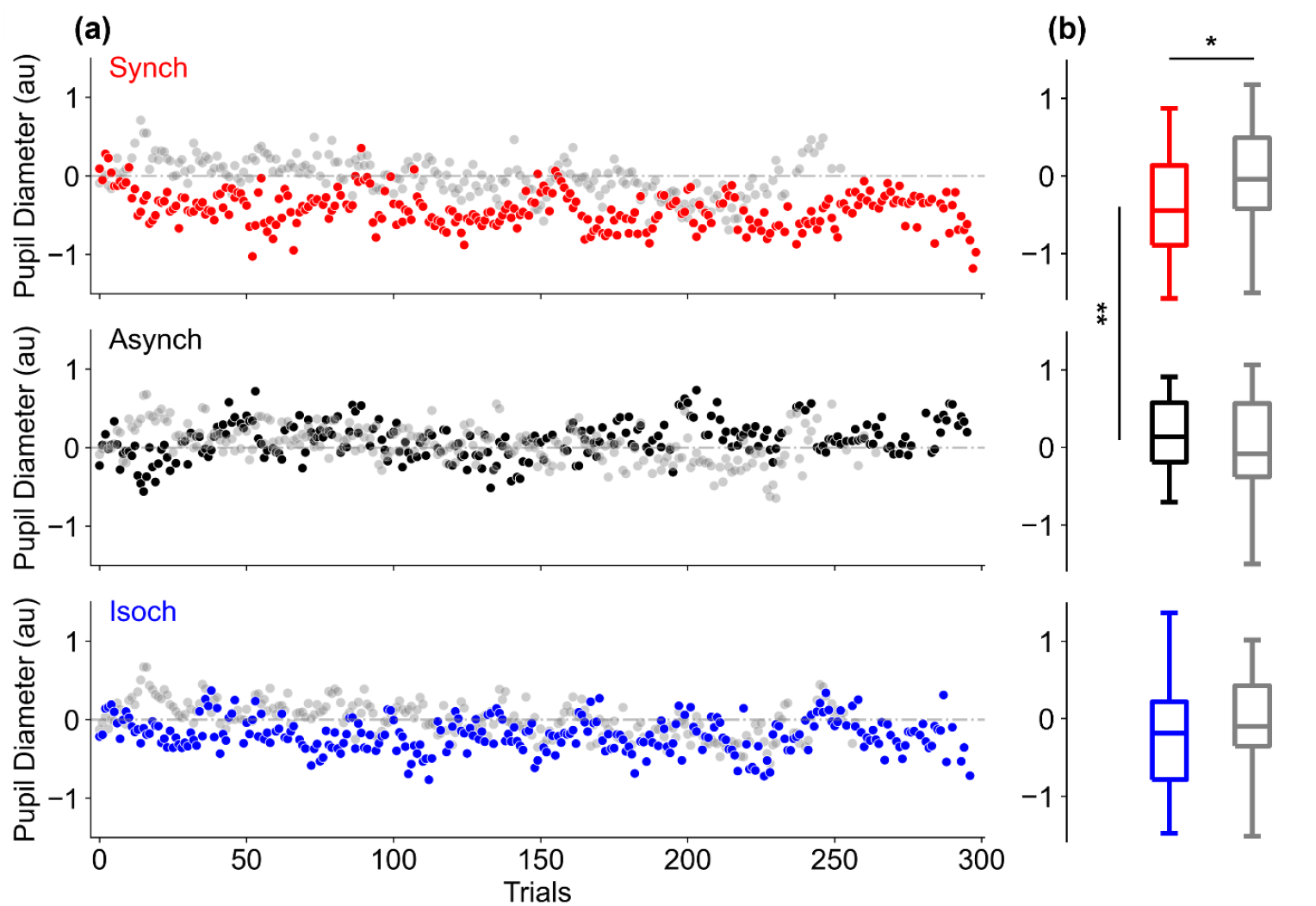
Global field power and pupil responses along the auditory regularity sequences. (a) Grand-averaged global field power (GFP) values over 300 trials for the Synch (red), Asynch (black), and Isoch (blue) auditory conditions (N = 31). Data are z-score normalized and referenced to the mean of the first 10 trials for each condition. (b) Distribution of GFP values averaged across the full sequence of 300 trials for each condition. (c) Grand-averaged pupil responses (N = 25; dashed lines) and GFP (N = 31; dotted lines) over 300 trials for each condition. Solid lines indicate best-fitting model (linear or exponential) selected based on the Akaike Information Criterion. In the Synch and Isoch conditions, both signals were best described by an exponential decay function. In the Synch condition, decay time constants were highly similar for pupil and GFP models (τ = 11.44 trials for pupil, τ = 13.30 trials for GFP), while in the Isoch decay rates diverged (τ = 6.88 for pupil, τ = 2.36 for GFP). In the asynch condition, both signals followed linear trends. Shaded regions represent ± standard error of the mean. au = arbitrary units; **p < 0.01 by two-tailed Wilcoxon signed-ranked tests.

To statistically compare this global pupil response across the auditory sequence, we calculated the average normalized pupil diameter over the 300 trials for each condition (Figure 4b). A Friedman test comparing global pupil diameter across the three auditory conditions revealed significant differences (W = 0.19, p < 0.01). Post-hoc Wilcoxon signed-rank tests indicated significantly higher constriction in the synch compared to the asynch condition (Kendall W = 66, p < 0.01), suggesting that cardio-audio predictability enhanced sustained pupil constriction. We also compared each auditory condition to its corresponding baseline. A significant difference was observed only in the synch compared to its baseline (W = 88, p < 0.05), further supporting a unique effect of heartbeat-locked auditory regularity on global pupil dynamics.

In parallel to the pupil analysis, GFP values were z-score normalized and referenced to the mean of the first 10 trials for each participant and condition (N = 31), allowing for the examination of neural activity changes relative to the beginning of each auditory sequence (Figure 5a). A Friedman test on the average GFP values across the 300 sounds revealed significant differences between auditory conditions (Figure 5b; Kendall W = 0.27, p < 0.001). Post-hoc Wilcoxon signed-rank tests showed that the synch condition elicited significantly higher GFP values compared to the asynch (W = 42, p < 0.001) and isoch (W = 106, p = 0.01) conditions. No significant difference was observed between the asynch and isoch conditions (W = 168, p = 0.19). This neural activity pattern mirrors the enhanced pupil constriction observed specifically in the synch condition, suggesting that cardio-audio regularity encoding uniquely modulated both autonomic and neural responses.

**Figure 5.**
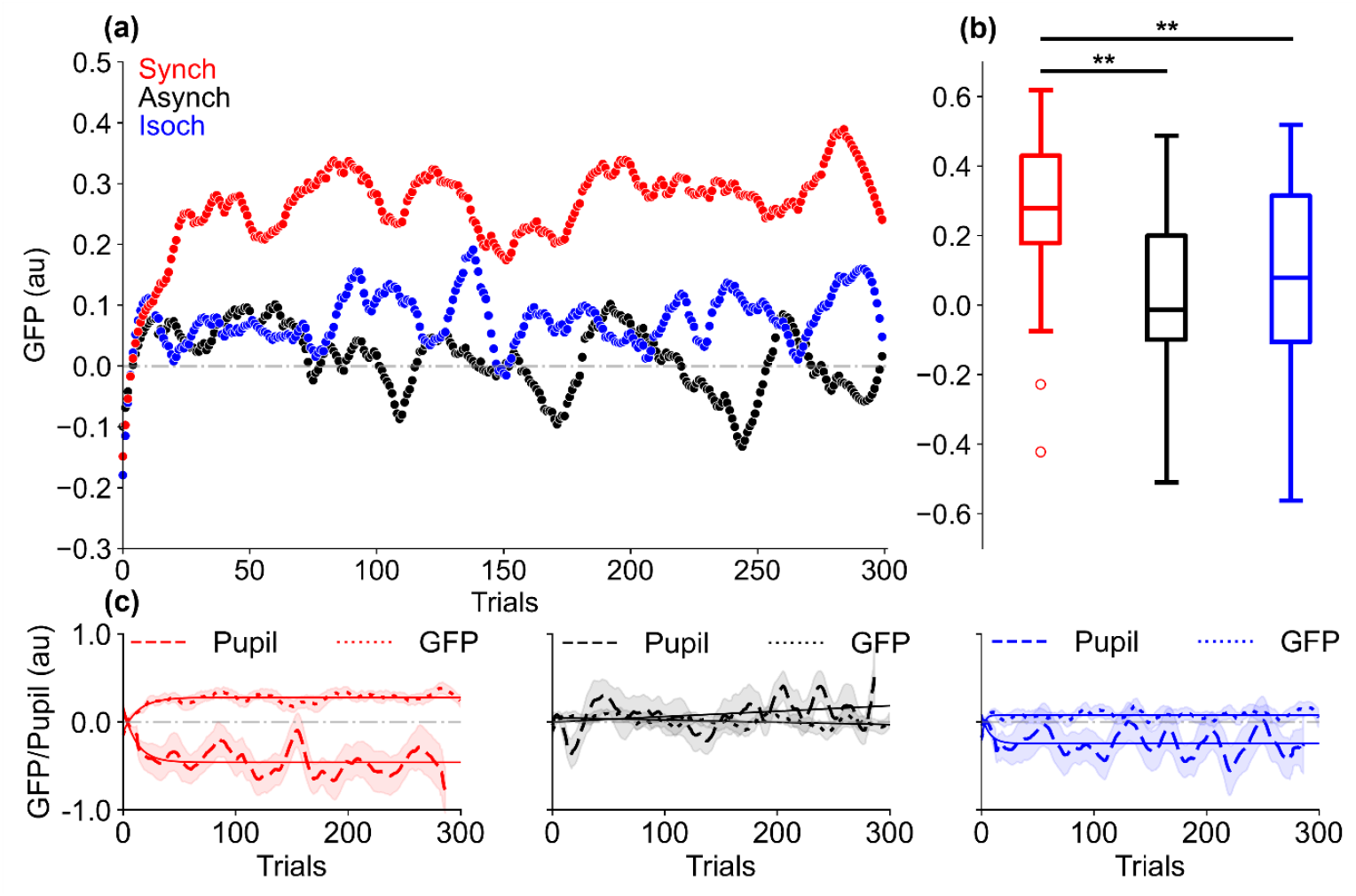
Global field power and pupil responses along the auditory regularity sequences. (a) Grand-averaged global field power (GFP) values over 300 trials for the Synch (red), Asynch (black), and Isoch (blue) auditory conditions (N = 31). Data are z-score normalized and referenced to the mean of the first 10 trials for each condition. (b) Distribution of GFP values averaged across the full sequence of 300 trials for each condition. (c) Grand-averaged pupil responses (N = 25; dashed lines) and GFP (N = 31; dotted lines) over 300 trials for each condition. Solid lines indicate best-fitting model (linear or exponential) selected based on the Akaike Information Criterion. In the Synch and Isoch conditions, both signals were best described by an exponential decay function. In the Synch condition, decay time constants were highly similar for pupil and GFP models (τ = 11.44 trials for pupil, τ = 13.30 trials for GFP), while in the Isoch decay rates diverged (τ = 6.88 for pupil, τ = 2.36 for GFP). In the asynch condition, both signals followed linear trends. Shaded regions represent ± standard error of the mean. au = arbitrary units; **p < 0.01 by two-tailed Wilcoxon signed-ranked tests.

To further explore this coupling, we modelled the temporal evolution of pupil diameter and GFP across the 300-trial auditory sequences (Figure 5c). Model selection using the Akaike Information Criterion (AIC)^30^ identified condition-specific dynamics. For the synch condition, exponential decay models (*y* = *a* · *e*^−*x/τ*^ + *c*) provided substantially better fits than linear models (*y* = *m* · *x* + *q)* for pupil responses (exponential AIC = -540.25 vs linear AIC = -510.98) and GFP (exponential AIC = -824.47 vs linear AIC = -725.92). Notably, the decay parameters were remarkably similar between pupil diameter (τ = 11.44) and GFP (τ = 13.30), indicating a tight temporal coupling between neural and autonomic responses. In the isoch condition, both measures also followed exponential decay patterns, although AIC differences were smaller, particularly for pupil responses (exponential AIC = -538.99 vs linear AIC = -535.70) compared to GFP (exponential AIC = -834.75 vs linear AIC = -817.08). However, the decay parameters differed substantially (τ = 6.88 for pupil and τ = 2.36 for GFP), suggesting weaker coupling between pupil and neural changes compared to the synch condition. In contrast, the asynch condition was best described by linear models for both pupil diameter (linear AIC = -500.13 vs exponential AIC = -498.14) and GFP (linear AIC = -789.86 vs exponential AIC = -764.78), reflecting a stable, non-adaptive temporal response profile. Together, these findings suggest that cardio-audio regularity encoding did not only modulate the magnitude of both neural and autonomic responses but also coordinated their temporal dynamics. The strong alignment observed in the synch condition points to a specific mechanism through which internal bodily rhythms can shape sustained sensory processing in both the brain as well as peripheral physiology.

### Relationship between pupil diameter and cardiac activity

To assess the link between tonic arousal, indexed by the heart rate, and overall pupil size during auditory stimulation, we correlated the global pupil response (averaged over the 300 sound trials with the average heart rate (beats per minute) across participants (N = 26), separately for each auditory condition (Figure 6a). A Shepherd’s Pi test revealed a statistically significant negative correlation in the synch condition only (r = -0.51 p = 0.01), pointing to an inherent relationship between pupil dilation and cardiac signal modulation in response to the heartbeat-locked auditory sequence. This negative correlation suggested that participants with slower heart rates exhibited greater pupil dilation than those with faster heart rates. To qualitatively verify this effect, we examined the time course of pupil diameter in two participant subgroups: those with the slowest (N = 10) and fastest (N = 10) heart rates (Figure 6b).

**Figure 6.**
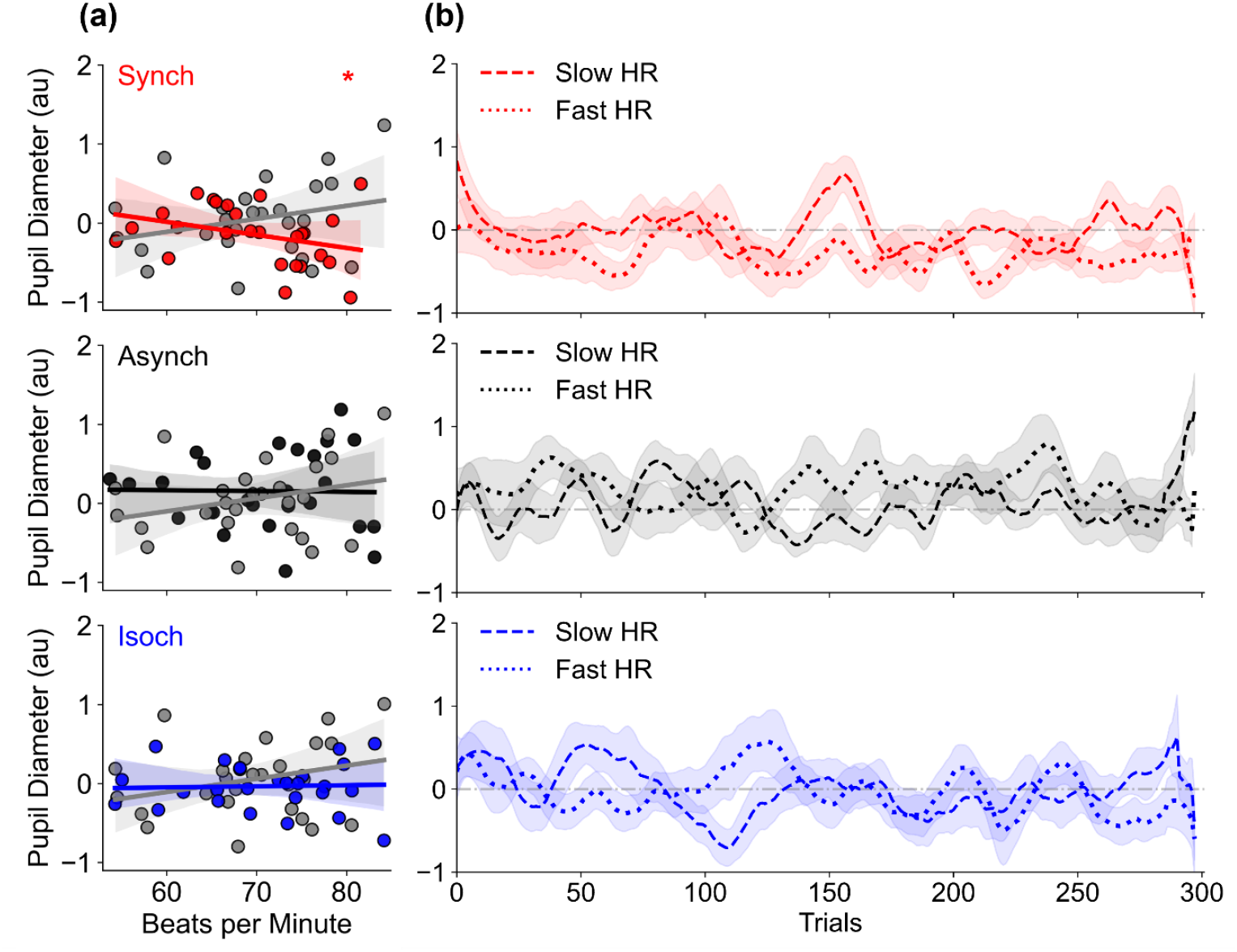
Relationship between pupil and cardiac activity across auditory regularity sequences. (a) Correlation between mean heart rate (beats per minute) and mean normalized pupil diameter across the 300-sound sequence for the Synch (red), Asynch (black), and Isoch (blue) conditions (N = 26). Corresponding values from the silent condition without auditory stimulation are shown in gray. (b) Grand-averaged time courses of pupil diameter (averaged over each trial window of -100 ms and 600 ms relative to sound onset), plotted for two subgroups of participants: those with the slowest (N = 10, dashed line) and fastest (N = 10, dotted line) heart rates (HR). Shaded regions represent ± standard error of the mean. au = arbitrary units; *p < 0.05 by Shepherd’s Pi correlation test.

## Discussion

We investigated the pupil and neural correlates of cardio-audio regularity encoding in 31 healthy participants by presenting sounds either in synchrony with the ongoing heartbeat (synch), at a fixed temporal pace (isoch), or with irregular pace (asynch), while keeping the average SS intervals matched across conditions. All auditory conditions elicited a pupil dilation following each sound that was different from baseline (no auditory stimulation). Similarly, all conditions evoked a typical auditory neural response, including a fronto-temporal N100 component. Uniquely in the synch condition, the AEP also showed an evoked response to the heartbeat due to the experimentally imposed relationship between heartbeat and sound. We then examined the global modulation of pupil and neural dynamics over 300-sound sequences. In the synch condition alone, we observed both a progressive pupil constriction and an increase in GFP relative to the beginning of the sequence. The pupil constriction was significantly higher compared to the asynch condition, and the GFP increase was significantly higher compared to both asynch and isoch conditions. Finally, we found a condition-specific relationship between average pupil diameter and heart rate. Specifically, in the synch condition, participants with faster heart rates showed higher pupil constriction compared to those with slower heart rate.

### Pupil response over the sequence

In line with previous studies, each auditory stimulus elicited a characteristic pupil response, consisting of an early constriction followed by a later dilation, observable within the first 500 ms after stimulus onset^12^ (Figure 2a). This response did not significantly differ across conditions, suggesting that short-latency pupil responses were not sensitive to the type of temporal regularity. In contrast, pupil dynamics over the entire 300-sound sequences revealed condition-specific effects (Figure 4a). Specifically, in the synch condition only, pupil dilation decreased later in the sequence compared to its beginning. This reduction was significantly different compared to both the asynch and baseline conditions (Figure 4b). This reduction suggests that the temporal alignment between auditory and cardiac signals, rather than regularity in SS intervals alone, modulates sustained autonomic responses. While a similar trend was observed in the isoch condition, it did not reach statistical significance.

Pupil diameter is widely used as a proxy for LC-NE activity^31–33,34,9,16^, which increases during arousing or information-rich (e.g., unexpected) events^10,11,8^. Such responses are believed to reflect widespread NE release from the LC, facilitating heightened information processing^35^. Along this line, the pupil constriction observed over the course of the synch sequence in our results suggests a decrease in LC-NE signals relative to the beginning of the sequence. While the start of the synch sequence presentation is characterized by high information content due to the inherent variability in SS intervals, it later shifts to a stable phase of high predictability, and therefore low information content, following the encoding of the cardiac and auditory temporal contingency. The encoding of the cardio-audio regularity would thus facilitate the anticipation of upcoming stimuli, reflected in the adaptation of pupil dilation over time^7,36^. Of note, while we expected similar constriction in the isoch condition due to the encoding of the temporal regularity (fixed SS intervals), this effect may have been too subtle, possibly because it involved only external auditory cues, whereas the synch condition leveraged both internal (cardiac) and external (auditory) signals. Thus, multimodal integration likely enhanced regularity encoding and the resulting autonomic response.

From a predictive coding perspective, NE has been proposed as a reset signal triggered by unexpected events and allowing for predictive model updates^12,8^. Pupil constriction during the synch condition may therefore reflect the sharpening of the predictive model priors, once the cardio-audio regularity is inferred. This observation suggests that pupil constriction mirrors the increase in signal predictability and the reduction in the degree of required model updating based on the incoming sensory evidence^7^. In our results, the integration of cardiac and auditory stimuli can be interpreted as an emergence of regularity that facilitates the processing of upcoming stimuli, resulting in pupil adaptation relative to the beginning of the sequence^7^. The constriction of the pupil diameter along the synch sequence may additionally relate to the modulation of arousal in the presence of regular vs random auditory sequences. The 300-stimulus presentation of the synch sequence, wherein a transition from an unpredictable to a predictable pattern of cardio-audio regularity is taking place, may result in reduced arousal and thus pupil constriction over the course of the sequence, possibly driven by facilitated sensory processing in the presence of highly predictable stimuli^12,36^.

### GFP increase during cardio-audio sequence presentation

At the neural level, average AEPs and GFP to individual sounds exhibited stereotypical components including the N100 topographical signature at 100 ms post-stimulus onset (Figure 3). As expected, these average responses were similar between the isoch and asynch conditions^24^, while in the synch, the neural response to the heartbeat signal shortly preceding the sound elicited a prominent voltage deflection just before the sound onset. These results show that, by construction and differently from the asynch and isoch, the synch sequence elicited a combination of two evoked responses to the cardiac and auditory input. Importantly, across the auditory sequence, the GFP increased significantly over time in the synch condition only (Figure 5a). This increase was significantly higher compared to both the isoch and asynch conditions (Figure 5b). Notably, in the synch, pupil constriction and GFP increase were best fit by exponential functions, with nearly identical time constants (∼12 trials), suggesting a tight coupling between the neural and autonomic systems under cardio-audio regularity (Figure 5c). In the isoch condition, both signals also followed exponential trajectories, but with different time constants (7 trials for pupil; 2 trials for GFP), suggesting that the encoding of purely external auditory regularity unfolded over a faster timescale compared to cardio-audio regularity inference in the synch condition. In contrast, asynch sequences followed linear trends (with no pupil constriction or GFP increase over time), consistent with lack of predictability.

This result aligns with previous findings of sustained GFP responses to extended auditory regular sequences, where increased GFP was similarly observed for regular compared to random sequences^19,21,37,38^. Within a predictive coding framework, such an increase has been suggested to reflect enhanced precision encoding in cortical regions as the brain generates a stable model of the sensory environment^19,39^. The GFP increase was higher in the synch condition as the priors were likely incorporating both the prediction of the cardiac and the auditory stimuli in contrast to the priors formed for the regularity established only between auditory stimuli in the isoch.

### Relationship between pupil and cardiac dynamics

We further examined the relationship between pupil diameter and heart rate at the single-subject level (Figure 6a). In line with our previous results, only in the synch sequence, we found a significant negative correlation: participants with faster heart rates showed reduced pupil dilation. As a result, the pupil constriction during the synch sequence was mostly likely driven by participants with faster heart rate (Figure 6b). Since faster heart rates led to shorter SS intervals, this result suggests that reduced arousal in these participants was tied to higher predictability from denser cardio-audio pairings. This observation is coherent with previous literature showing that longer SS intervals evoke greater arousal, as indexed by galvanic skin responses^40,41^. This relationship was absent in the isoch and asynch conditions, supporting the notion that only heartbeat-synchronized auditory inputs modulate the pupil-heart coupling (Figure 4).

### Limitations

Our study had several limitations. First, the relatively short SS intervals limited our ability to track pupil dynamics over longer timescales, which may evolve over seconds. Thus, potential modulations induced by auditory regularities occurring later than the average SS interval may have been overlooked. Second, participants were not engaged in any task, which limited control over attention and task engagement, particularly in a prolonged experimental session (∼2-3 hours), potentially weakening pupil and/or neural modulation effects. Third, while our design enabled the investigation of sustained regularity encoding using pupil and neural measures, future studies should incorporate deviant stimuli or violations within the sequence to assess how prediction errors influence pupil and neural dynamics.

### Conclusion

In summary, our findings extend previous work on cardio-audio regularity encoding^23–25^, showing that this process influences both neural and autonomic (pupil diameter) systems dynamics. Heartbeat-synchronized sounds enhanced predictive processing, as indicated by reduced arousal (pupil adaptation) and enhanced precision (GFP increase). These effects were specific to cardio-audio regularity, suggesting that the integration of internal bodily rhythms with external sensory inputs facilitates sequence regularity encoding and sustained sensory prediction.

## Materials and methods

### Participants

Thirty-one right-handed healthy volunteers (16 female, mean ± standard deviation (SD) of age = 23.42 ± 3.40) were recruited for this study as part of an experiment investigating auditory regularity processing in wakefulness and sleep. A semi-structured interview confirmed the absence of psychiatric, neurological, respiratory or cardiovascular conditions. Participants had normal hearing, were non-smokers and did not take any psychoactive medication. A regular sleep schedule without excessive daytime sleepiness or sleep disturbances was confirmed using the Pittsburgh Sleep Quality Index and the Epworth Sleepiness scale, respectively. Approval for the study (Project-ID: 2020-02373) was obtained by the local ethics committee (La Commission Cantonale d’Ethique de la Recherche sur l’Etre Humain), in accordance with the Helsinki declaration. All participants gave written informed consent as per the ethics agreement and were naive to the experimental manipulation implemented in the study, verified by an informal interview following the completion of the experiment.

### Experimental procedures

This study followed a two-way crossover experimental design wherein participants performed three consecutive overnight sleep sessions and a separate wakefulness session, the latter taking place up to a week before the first sleep session or up to a week after the last sleep session (wakefulness session was first in 15 participants). Acquired data and procedures were matched between wakefulness and sleep, with the exception of the eye-tracking data acquisition, only feasible in wakefulness. The study outlined herein utilized only data during wakefulness hence, only experimental procedures for wake are described.

During the wakefulness session, participants performed an auditory stimulation experiment in a dimly lit experimental booth isolated from sound and electrical interference. They sat on a chair with the head fixed using the eye-tracker chin rest, placed approximately 55 cm from the screen. Ocular data were acquired at 500 Hz using the SR Eyelink 1000 plus (SR Research Ltd., Mississauga, Canada). 64-channel (waveguard net, eego mylab, ANT Neuro, Hengelo, Netherlands) EEG, ECG, horizontal and vertical electrooculography, submental electromyography, and respiration (via an upper abdominal respiratory belt) were acquired at 1024 Hz using the ANT Neuro system (eego mylab, ANT Neuro, Hengelo, Netherlands) and lab streaming layer (LSL^42^, https://github.com/sccn/labstreaminglayer). Following electrophysiological signals’ preparation, calibration of the right eye was performed. In five participants, the right eye could not be successfully tracked and therefore, the left eye was used. A central white cross was displayed on a gray background for the remainder of the experiment which participants were asked to fixate on. Prior to data acquisition, the accurate real-time detection of R peaks in the ECG signal was ensured by tuning the height and prominence parameters as part of the python function findpeaks.py, which was used to select R peaks during the experiment.

Sounds were 1000 Hz sinusoidal 16-bit stereo tones of 100 ms duration and 0 μs inter-aural time difference sampled at 44.1 KHz and generated using the sound function in Psychopy^43^. A tukey window was applied to each tone. Sound stimuli were presented binaurally through in-earphones with an intensity adjusted at the single-subject level to a comfortable level, determined prior to the experimental paradigm administration.

Paradigm administration was performed using Python-based Psychopy and custom-made Python scripts. The experimental paradigm included the administration of three types of auditory sequences with 300 sound stimuli each and a condition without auditory stimulation (herein referred to as ‘silent’ or ‘no auditory stimulation condition’). The silent condition lasted 4 minutes. In the synch auditory condition, sounds were presented at a fixed 52 ms R peak-to-sound (RS) interval after the online detection of an R peak in the ECG signal. In the isoch auditory condition, the median RR interval from a preceding synch block served as the fixed SS interval. Finally, in the asynch auditory condition, the RR intervals from a preceding synch block were shuffled and served as the SS intervals, following exclusion of extreme RR interval outliers. Outliers were identified if they were above 1.8 times or below 0.2 times the mean SS interval in the synch condition. A series of control analyses verified the correct administration of the experimental paradigm (Figure S6). First, experimental blocks with frequent flawed R peak real-time detection or faulty auditory stimulus presentation were excluded from a given participant’s dataset. Second, Friedman tests (p < 0.001) with within-subject factor auditory condition (synch, asynch, isoch) and post-hoc Wilcoxon signed-rank tests with Bonferroni correction for multiple comparisons confirmed a fixed RS interval of 52 ms with a variability (standard deviation (SD) of RS intervals here and elsewhere) of 1 ms in the synch, significantly different (p < 0.001) to the asynch and isoch conditions with intervals of -3 ms and -2 ms and variabilities of 262 ms and 259 ms, respectively (Figure S6a). In addition, we observed no variability in SS intervals (SD of SS intervals = 0 ms) in the isoch condition, significantly different (p < 0.001) to the synch and asynch condition with values of 77 ms and 75 ms, respectively (Figure S6b). Finally, as expected no statistically significant differences (p > 0.05) were observed in RR intervals and variabilities (Figure S6c) across all experimental conditions (synch, asynch, isoch, silent).

During wakefulness, experimental conditions were presented in blocks of four, followed by a pause to allow for the participant to rest. The first block always began with a silent condition followed by a synch condition to generate the sequence of SS intervals in the isoch and asynch conditions, which were presented in a randomized order. All other blocks had a randomized order presentation with the only rule being that a block of synch, asynch, isoch and silent were presented before a pause. All conditions within a block were separated by an interval of 30 seconds and the experimental block of four conditions lasted approximately 20 minutes (depending on the individual’s heart rate). A total of six experimental blocks were administered. The duration of each block was comparable to previous experiments examining pupil responses to auditory regularities (e.g., up to 288 trials with 2-3 s stimuli and comparable SS intervals in passive and active tasks^11^; 56-minute active task with 168 trials^12^; 26 sounds repeated up to 12 times in an active task^7^; 20-minute passive listening experiment^29^).

### Data analysis

Data analyses were performed in Python 3.12 and MATLAB (R2019b, The MathWorks, Natick, MA) using open-source toolboxes MNE-python^44^, EEGLAB (13.4.4b)^45^, Fieldtrip (20231015)^46^ and custom-made scripts.

### Pupil data analysis

Blinks in ocular data were detected in real-time by the eye-tracking software (SR Research Ltd., Mississauga, Canada). In addition, blinks identified as missing data of maximum 300 ms were interpolated using MNE’s *interpolate_blinks* function.

#### Pupil data preprocessing

Pupil data were segmented in trials of 700 ms, between -100 ms and 600 ms from each sound onset for each auditory condition (synch, asynch, isoch). For the silent condition in the absence of auditory stimulation, we defined ‘artificial’ sound onsets by matching the SS intervals to those in the asynch and isoch conditions. For the synch condition, due to fixed relationship between R peak and sound, ‘artificial’ sound onsets were placed 52 ms after each R peak in the silent condition, based on the average RS delay observed across participants in the synch. Pupil data during the silent condition were segmented in trials of 700 ms, between –100 ms and 600 ms, to generate a baseline dataset for each auditory condition.

To investigate the impact of contiguous auditory stimuli on pupil diameter, we additionally performed a multi-trial analysis for triplets of sounds. This analysis was only performed in the isoch condition since fixed SS intervals during contiguous trials were essential to generate reliable trial averages. The dataset was first segmented around triplets of contiguous trials with a trial length of 2200 ms, between -100 ms after the first sound onset and 600 ms after the third sound onset. The SS interval was calculated for each trial (mean of the two SS intervals in a triplet) and if the mean was not equal to the SS of the whole condition, the trial was rejected. Next, the trials with the highest number of matched SS intervals were retained, which included participants with SS intervals between 800 ms and 900 ms.

Upcoming pre-processing steps were performed for the pupil data in response to single sounds and triplets of sounds. As a first step, we removed trials with overlapping data to the preceding trial. Next, artifacted trials were excluded from further analysis following standard procedures^47^ as follows. First, trials containing any missing values not previously identified as blinks, were discarded. Second, to remove trials with unrealistic pupil diameter values, trials with one or more data points exceeding the mean ± 3SD (± 5SD for multi-trial analysis) were rejected, where the mean and SD was computed along the trial length. Third, we excluded trials with a rapid and therefore non-physiological pupil diameter change by evaluating the mean and SD of the pupil signal over moving windows of 60 ms along the trial length. Trials with *≥* 3 points (*≥* 5 for multi-trial analysis) exceeding the mean ± 3SD within at least one 60 ms window were rejected. Fourth, for each subject, block, and auditory condition or baseline, the mean and SD were calculated to identify outlier trials. Trials with one or more samples exceeding the mean ± 3SD (± 5SD for multi-trial analysis) were removed. Analyses were conducted within each of the six experimental blocks and across all six blocks. For the within-block analysis, participants were included if they had a minimum of 100 trials for each condition. For the across-blocks analysis, participants were eligible if they were included in a minimum of three blocks for the within-block analysis.

After the aforementioned pre-processing steps, 221 ± 44 and 222 ± 75 (mean ± SD) trials were retained for the analyses of the pupil response to single sound and to triplets of sound, respectively. Finally, for each participant, pupil data were z-score normalized within each experimental block, making them comparable across the volunteer cohort.

#### Pupil response to sounds

The local effect of single sounds on pupil diameter was performed for each experimental condition (synch, asynch, isoch, baseline) and the multi-trial analysis on the effect of triplets of sounds was performed only in the isoch condition. When analyzing sound local effects, each sound trial was referenced to the median pupil value of the 100 ms pre-stimulus period. Non-parametric repeated-measures Friedman tests (p < 0.05) with within-subject factor auditory condition (synch, asynch, isoch) and post-hoc one-tailed Wilcoxon signed-rank tests (p < 0.05) assessed whether significant differences in local effects existed in the average local pupil response across timepoints in a trial. In addition, one-tailed Wilcoxon signed-rank tests (p < 0.05) contrasted each auditory condition to its corresponding baseline.

#### Pupil responses to sound sequences

To investigate the modulation of the pupil diameter over sustained sound exposure within each sequence of 300 sounds from a given experimental block, we computed the normalized pupil diameter averaged over the course of the single-trial (between -100 ms and 600 ms without referencing to the -100 ms to 0 ms pre-stimulus period) for each administered sound (synch, asynch, isoch) or ‘artificial’ sound (baseline). Each data point corresponded to the ordinal sequence of the administered sound such that the *nth* data point corresponded to the average pupil response of the *nth* sound administered within the sequence. Of note, data points for each ordinal position within the sequences were not present for all participants following pupil data pre-processing, yielding a variable number of participants contributing to the grand-average global effect for each data point. To address this, we required a minimum of 15 contributing participants for each data point, which resulted in the inclusion of 202 ± 38 data points for each experimental condition and participant.

Next, for each participant, we computed the mean of the first 10 trials in each block and condition, which was used as a reference for pupil dilation at the beginning of the sequence and subtracted from all trials to highlight changes over repeated auditory stimulation within each sequence of 300 sounds. Only participants who had at least 1 trial present in the first 10 trials were included in this analysis. Next, to statistically compare the global effect, we computed the average pupil dilation over all data points in the sequence of 300 sounds or ‘artificial’ sounds at the single-subject level. Friedman tests (p < 0.05) with within-subject factor auditory condition (synch, asynch, isoch) and post-hoc two-tailed Wilcoxon signed-rank tests (p < 0.05) assessed whether significant differences existed across conditions. In addition, two-tailed Wilcoxon signed-rank tests (p < 0.05) contrasted each auditory condition to its corresponding baseline.

### EEG data analysis

Raw EEG data were band-pass filtered using second-order Butterworth filters between 0.5 and 30 Hz. Bad channels were excluded, and independent component analysis was used to identify and remove any cardiac field, eye or muscle related components. Next, bad channels were interpolated using spherical splines^48^ with an average (± SD) of 7.3 ± 2.5 interpolated channels. EEG data were epoched between −100 ms and 600 ms relative to the sound onset and artifacted trials were identified using a semi-automated approach by utilizing the Fieldtrip function *ft_rejectvisual*, verified by the visual inspection of selected trials. An average (± SD) of 286 ± 4 AEP trials were retained within each block. Finally, EEG data were re-referenced to the common average and the grand-average AEPs were generated for each auditory condition (synch, asynch, isoch), experimental block and participant. In addition, the single-trial GFP was calculated as the standard deviation of the EEG magnitude across channels for each timepoint in a given trial. Finally, for each participant, GFP values were z-score normalized within each experimental block. Similar to the global effect for the pupil diameter, sustained GFP over the repetition of 300 sounds within each auditory condition block was referenced to the mean GFP of the first 10 trials.

### Mathematical modelling of the pupil diameter and GFP global effect

To characterize the temporal dynamics of the pupil and GFP responses during sustained auditory stimulation, mathematical modeling was applied to the global effect data across the 300 trials within a sequence in the three auditory conditions (synch, asynch, isoch). For each participant and condition, linear interpolation was performed to fill missing data points in the trial-by-trial sequences, followed by smoothing using a Savitzky-Golay filter^49^ (polynomial order = 3, window size = 25 samples). The smoothed single-subject curves were then averaged across participants to obtain grand-averaged temporal profiles for each condition. Two models were fit to the grand-averaged data: (i) an exponential decay model of the form *y* = *a* · *e*^−*x/τ*^ + *c*, where *a* represents the initial amplitude, τ is the decay parameter, and *c* is the asymptotic value; (ii) a linear model of the form *y* = *m* · *x* + *q*, where *m* is the slope and *q* is the intercept. Model selection was performed using the AIC, with the model yielding the lowest AIC value considered the best fit for each condition and signal type. For exponential models, the time constant τ was calculated as τ = -1/*b*, representing the characteristic decay time in trials.

### Pupil diameter and heart rate correlation analysis

To assess the relationship between normalized pupil responses and RR intervals, correlation analyses were conducted. R-peaks in the ECG were detected using the Pan and Tompkins algorithm^50^ and unrealistic RR intervals due to flawed offline R-peak detection were excluded using the *rmoutliers* function in MATLAB. For each participant and experimental block, we calculated the mean heart rate (in beats per minute) and mean normalized pupil diameter based on the average of RR intervals and averaged single-trial pupil diameter (between -100 ms and 600 ms ms without referencing to the -100 ms to 0 ms pre-stimulus period) for each administered sound (synch, asynch, isoch) or ‘artificial’ sound (baseline) over the sequence of 300 sounds. The correlation between these measures was then computed across participants, separately for each experimental condition. To account for potential outliers, the Shepherd’s Pi robust correlation (p < 0.01) was used^51^, which applies a non-parametric Spearman correlation after removing outliers identified through a bootstrap estimation of the Mahalanobis distance between samples.

To further examine the relationship between heart rate and pupil dynamics, participants were divided into slower and faster heart rate groups based on their median heart rate. To ensure consistent numbers of participants in each group and to maximize group separation, we implemented a two-step selection procedure. First, we considered the range of heart rate values across participants and conditions, and we separated these values in two groups based on whether they were below or above the median. We considered participants whose heart rate was always lower or higher than this median across conditions. Second, to establish balanced groups in the two heart rate ranges, we excluded additional participants from the higher heart rate group by removing those with the lowest heart rates, thereby increasing the inter-group heart rate difference. This process yielded two distinct groups, with each participant consistently represented across all auditory conditions. Finally, we qualitatively compared the two groups’ global pupil responses by looking at the averaged single-trial pupil diameter (between -100 ms and 600 ms without referencing to the -100 ms to 0 ms pre-stimulus period) for each administered sound (synch, asynch, isoch) over the sequence of 300 sounds.

## Data and code availability

Data can be made available by the corresponding author upon reasonable request.

The code for the experimental paradigm administration (https://github.com/fcbg-platforms/eeg-cardio-audio-sleep/tree/maint/0.3) and for the data analysis (https://github.com/GiovanniChiarion/CardioAudio_Pupil.git) are publicly available.

## Acknowledgements

The authors thank Matthieu Scheltienne from the Campus Biotech Foundation in Geneva for assistance with the design of the code for the experimental paradigm administration. In addition, the authors extend their gratitude to Gwenael Birot from the Campus Biotech Foundation in Geneva for useful feedback and guidance with regards to experimental procedures. Finally, we thank Alice Clerget and Erin Mahan for assistance with participant recruitment and data acquisition.

This work was supported by the Bertarelli Foundation grant ‘Catalyst’ to MDL and SS, the Swiss National Science Foundation (grant 32003B_212981) and the Eurostars project E!3489 to MDL, and the University of Lausanne ‘Transition Grant’ to AP.

## Author contributions

**Giovanni Chiarion:** Data curation, Methodology, Formal analysis, Writing - original draft; **Jacinthe Cataldi:** Data curation, Formal analysis, Writing - review and editing; **Sophie Schwartz:** Conceptualization, Resources, Funding acquisition, Project administration, Writing - review and editing; **Andria Pelentritou:** Conceptualization, Data acquisition, Data curation, Methodology, Formal analysis, Supervision, Writing - original draft; **Marzia De Lucia:** Conceptualization, Methodology, Funding acquisition, Project administration, Supervision, Writing - original draft.

