## Supplemental Information for "Cardiac-locked auditory stimulation modulates pupil and neural dynamics"

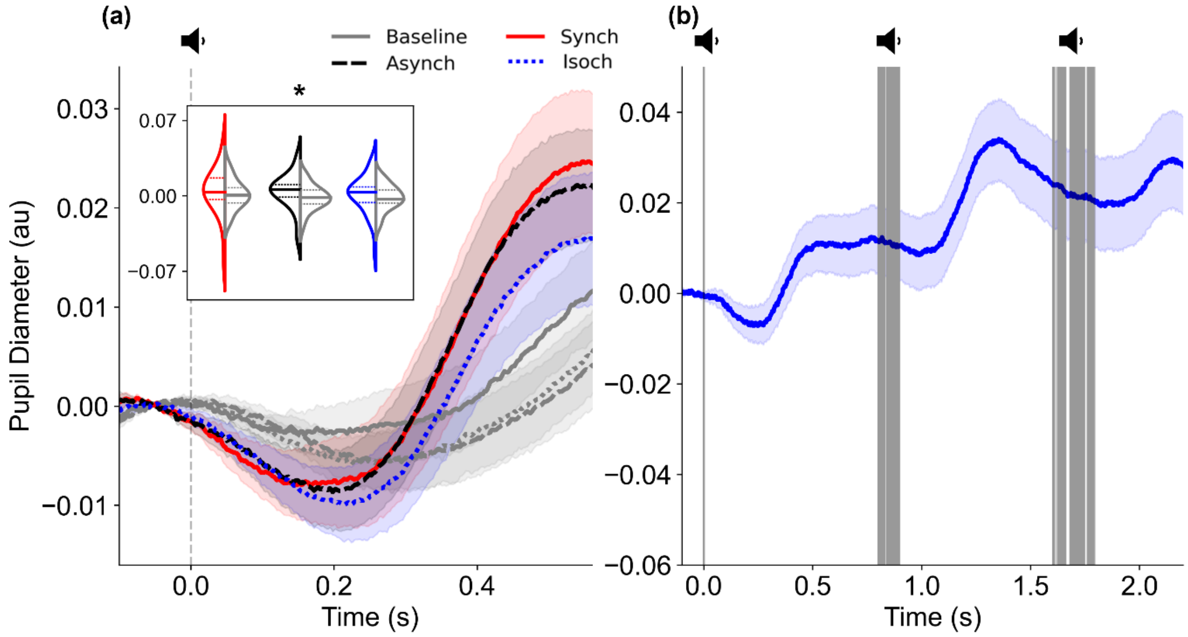


**Figure S1**. **Pupil responses to sounds across experimental blocks.** (a) Grand-averaged pupil diameter (N = 27) time-locked to sound onset (vertical dashed gray line) for the six experimental blocks across all conditions: Synch (red), Asynch (black), Isoch (blue), and their corresponding Baselines (gray). The Baseline responses represent pupil activity without sound presentation, epoched relative to 'artificial' sound onsets that matched the temporal structure of the auditory conditions. Pupil diameter data were z-score normalized and referenced to the median pupil diameter during the 100 ms pre-stimulus period. The inset shows violin plots of the averaged pupil diameter over the entire trial period (between -100 ms and 600 ms) for each auditory condition and its corresponding baseline. Horizontal lines within each violin plot indicate the first quartile, median, and third quartile. (b) Grand-averaged pupil diameter (N = 21) in response to sequences of three consecutive sounds (vertical gray lines) for the six experimental blocks in the Isoch condition, time-locked to the first sound. Data were z-score normalized and referenced to the median pupil diameter during the 100 ms pre-stimulus period of the first sound. Shaded areas in both panels represent ± standard error of the mean. au = arbitrary units; *$p < 0.05$ by one-tailed Wilcoxon signed-rank tests.

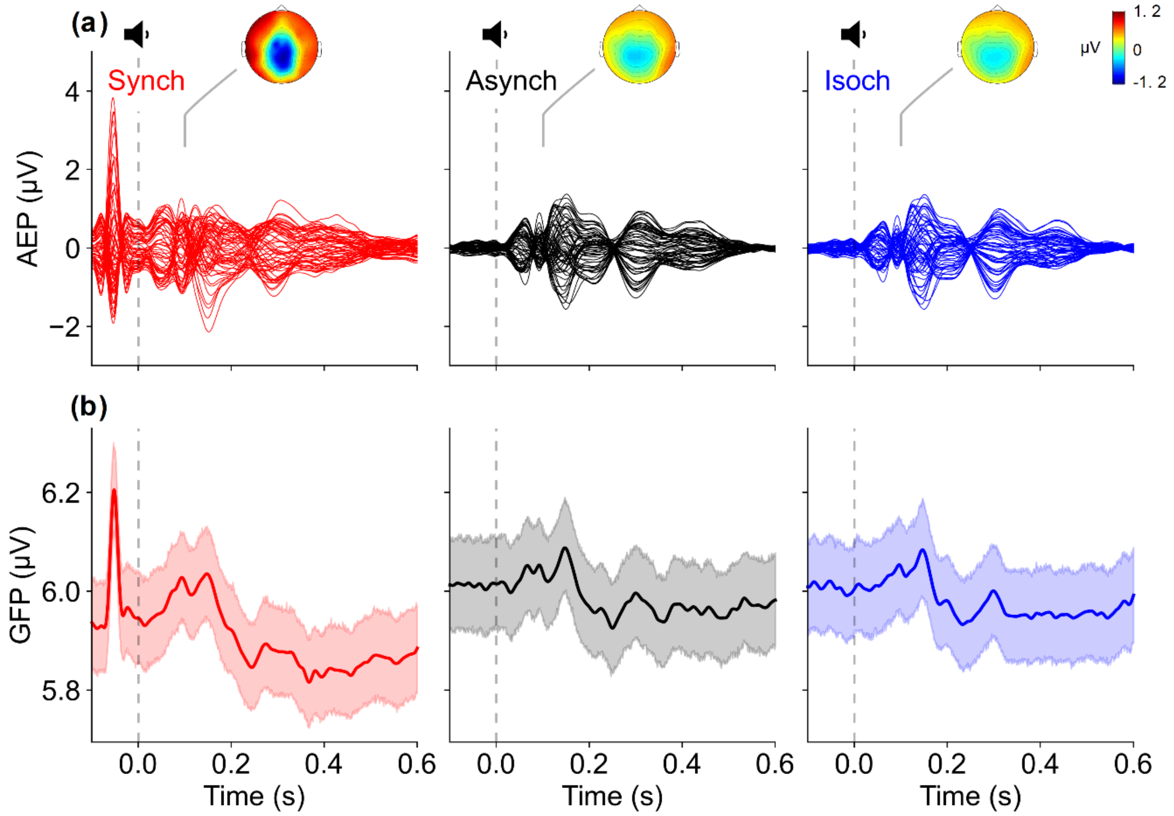


**Figure S2. Neural auditory evoked responses across experimental blocks.** (a) Grand-averaged auditory evoked potentials (AEPs; N = 31), time-locked to sound onset (vertical dashed gray line) for the six experimental blocks across the three auditory conditions: Synch (red), Asynch (black), and Isoch (blue). The topographic maps show the scalp distribution of AEP amplitudes at 100 ms post-stimulus onset. (b) Grand-averaged global field power (GFP) for the six experimental blocks across the three auditory conditions, representing the overall strength of neural activity across all electrodes over time. GFP was computed as the standard deviation of EEG potentials across all electrodes at each time point, providing a reference-independent measure of global brain activity. Shaded regions represent ± standard error of the mean.

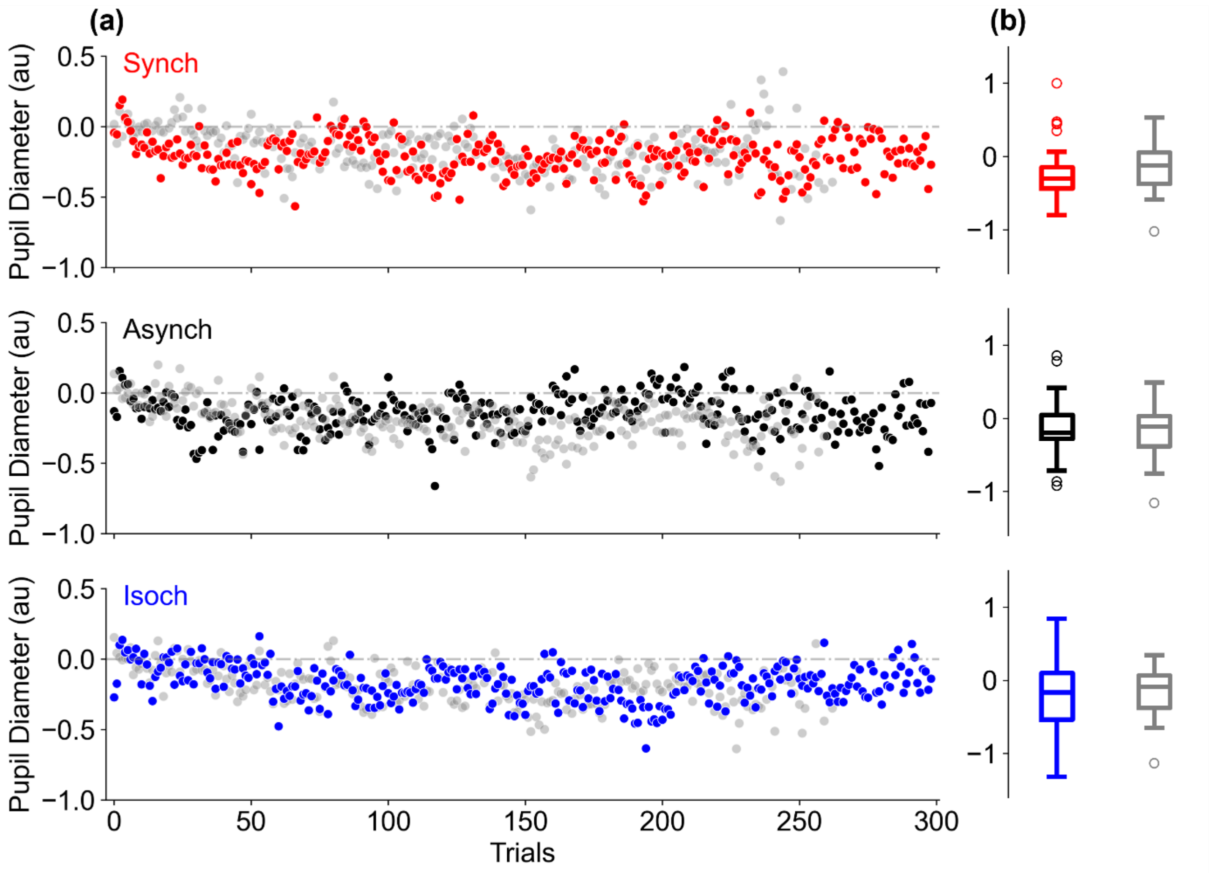


**Figure S3. Pupil responses along the auditory regularity sequences across experimental blocks.** (a) Normalized pupil diameter over the course of 300 auditory stimuli in the Synch (red), Asynch (black), and Isoch (blue) conditions, as well as 'artificial' onsets in their corresponding Baselines (grey) without auditory stimulation for the six experimental blocks. Data points represent the pupil response averaged across participants (N = 25) for each trial (between -100 ms and 600 ms time-locked to individual sounds), referenced to the mean of the first 10 trials for each condition. Negative values indicate pupil diameter constriction relative to the beginning of the sequence. (b) Boxplots showing the distribution of average pupil diameter over the entire sequence (300 sounds) for each auditory condition and its corresponding baseline. au = arbitrary units.

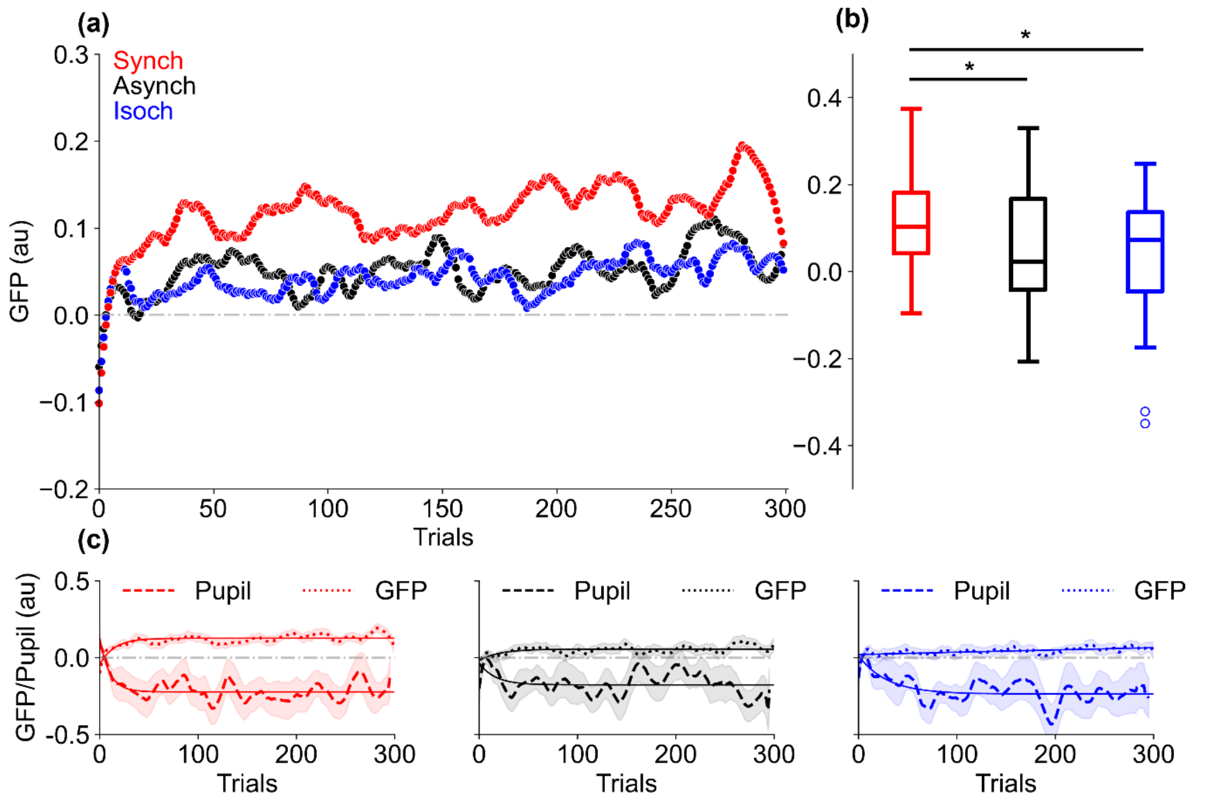


**Figure S4. Global field power and pupil responses along the auditory regularity sequences across experimental blocks.** (a) Grand-averaged global field power (GFP) values over 300 trials for the Synch (red), Asynch (black), and Isoch (blue) auditory conditions and for the six experimental blocks (N = 31). Data are z-score normalized and referenced to the mean of the first 10 trials for each condition. (b) Distribution of GFP values averaged across the full sequence of 300 trials for each condition. (c) Grand-averaged pupil responses (N = 27; dashed lines) and GFP (N = 31; dotted lines) over 300 trials for each condition and for the six experimental blocks. Solid lines indicate best-fitting linear or exponential models selected based on the Akaike Information Criterion. In the Synch condition, both signals were best described by an exponential decay function with similarly small time constants ($\tau = 9.98$ trials for pupil, $\tau = 14.62$ trials for GFP), indicating concordant temporal dynamics. In the Asynch condition, both signals were best described by exponential decay functions with higher time constants ($\tau = 16.17$ trials for pupil, $\tau = 19.86$ trials for GFP). In the Isoch condition, pupil responses followed an exponential decay ($\tau = 29.28$ trials) while GFP showed linear trends. Shaded regions represent ±standard error of the mean. au = arbitrary units; **$p < 0.05$ by two-tailed Wilcoxon signed-ranked tests.

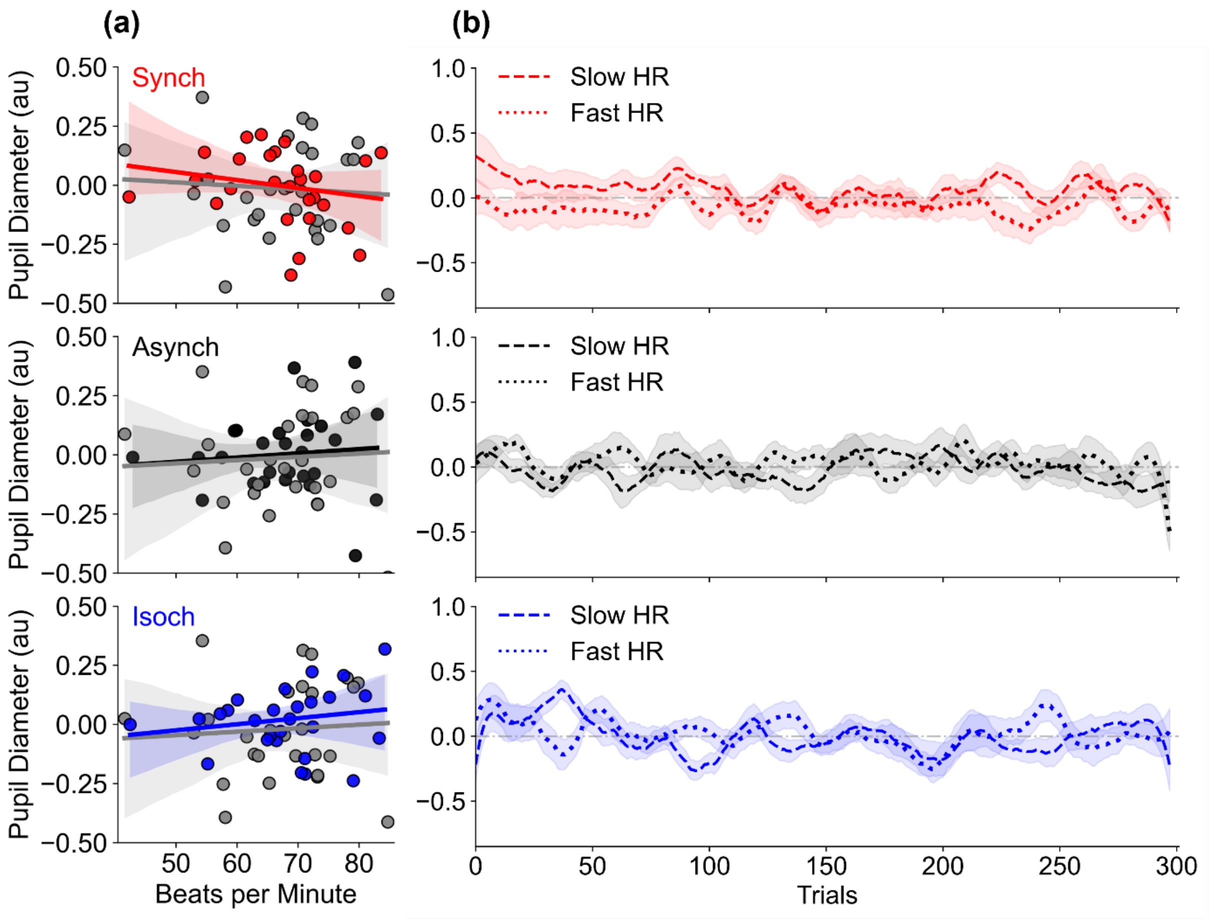


**Figure S5. Relationship between pupil and cardiac activity across auditory regularity sequences across experimental blocks.** (a) Correlation between mean heart rate (beats per minute) and mean normalized pupil diameter across the 300-sound sequence for the Synch (red), Asynch (black), and Isoch (blue) conditions for the six experimental blocks (N = 27). Corresponding values from the silent condition without auditory stimulation are shown in gray. Statistically significant correlations were identified using Shepherd's Pi test ($p < 0.05$). No condition showed a statistically significant correlation: synch ($r = -0.38$, $p = 0.06$), asynch ($r = 0.11$, $p = 0.61$), and isoch ($r = 0.19$, $p = 0.37$). Correlations during baseline were also not significant: synch baseline ($r = 0.31$, $p = 0.15$), asynch baseline ($r = 0.31$, $p = 0.14$), isoch baseline ($r = 0.33$, $p = 0.11$). (b) Grand-averaged time courses of pupil diameter for the six experimental blocks (averaged over each trial window of -100 ms and 600 ms relative to sound onset), plotted for two subgroups of participants: those with the slowest (N = 13, dashed line) and fastest (N = 13, dotted line) heart rates (HR). Shaded regions represent ± standard error of the mean. au = arbitrary units.

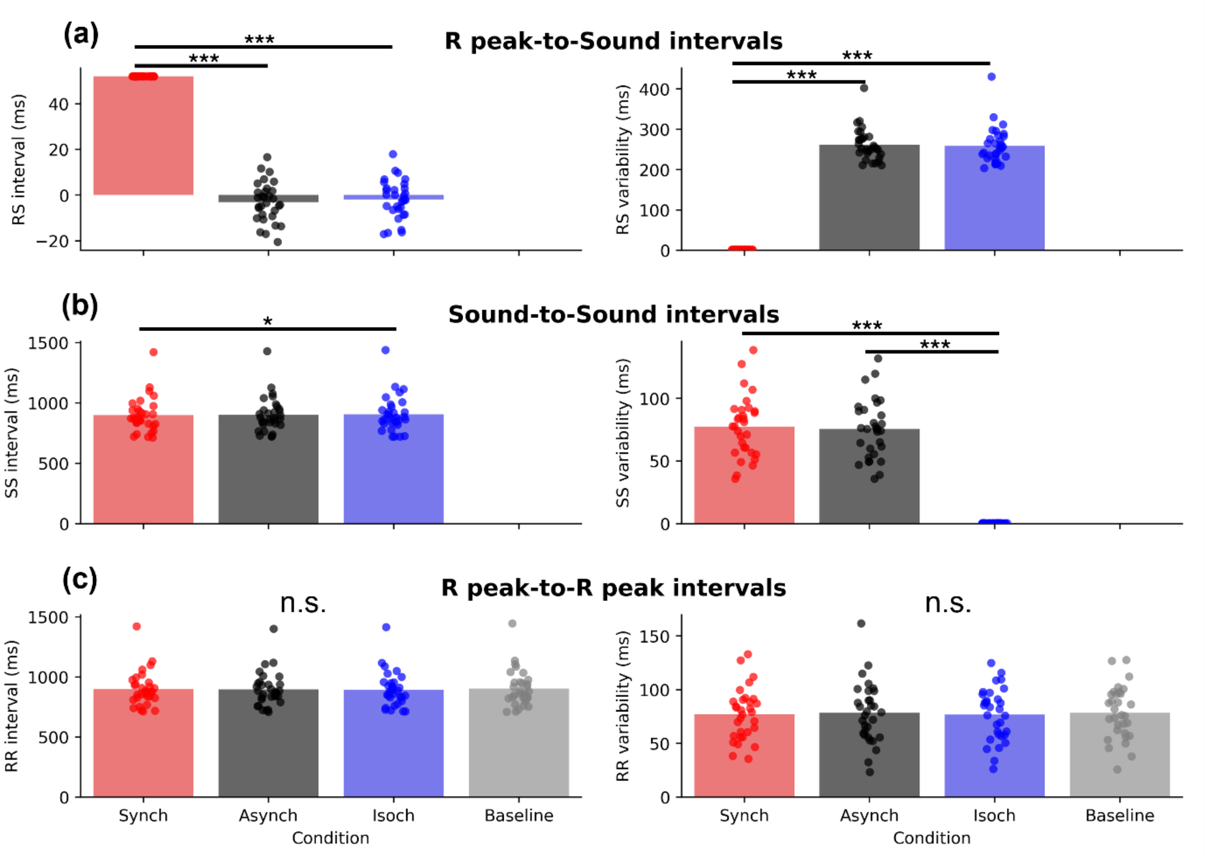


**Figure S6. Experimental paradigm control analysis.** (a) R peak-to-Sound (RS), (b) Sound-to-Sound (SS), (c) R peak-to-R peak (RR) intervals and variabilities (standard deviation of intervals) for the synch (red), asynch (black), isoch (blue), and baseline (gray, where applicable) conditions for the six experimental blocks (N = 31). Individual data points represent single-participant values. In the synch condition, RS variability was minimal, confirming a precise synchronization of cardiac and auditory stimuli. The isoch condition showed minimal SS variability while variable intervals were observed in the synch and asynch conditions. Finally, as expected, RR intervals and variabilities were matched across experimental conditions, and SS intervals were matched between the regular (synch, isoch) and irregular (sequence) across auditory conditions. ***$p < 0.0001$, *$p < 0.0160$, n.s. = not significant by Wilcoxon-signed rank tests with Bonferroni correction for multiple comparisons.
